# Elephant trunks use an adaptable prehensile grip

**DOI:** 10.1101/2022.10.06.511214

**Authors:** Andrew K. Schulz, Joy S. Reidenberg, Jia Ning Wu, Cheuk Ying Tang, Benjamin Seleb, Josh Mancebo, Nathan Elgart, David L. Hu

## Abstract

Elephants have long been observed to grip objects with their trunk, but little is known about how they adjust their strategy for different weights. In this study, we challenge a female African elephant at Zoo Atlanta to lift 20 to 60 kg barbell weights with only its trunk. We measure the trunk’s shape and wrinkle geometry from a frozen elephant trunk at the Smithsonian. We observe several strategies employed to accommodate heavier weights, including accelerating less, orienting the trunk more vertically, and wrapping the barbell with a greater trunk length. Mathematical models show that increasing barbell weights are associated with constant trunk tensile force and an increasing barbell-wrapping surface area due to the trunk’s wrinkles. Our findings may inspire the design of more adaptable soft robotic grippers that can improve grip using surface morphology such as wrinkles.

## 1 Introduction

In this study, we investigate the elephant trunk’s prehension, the ability to grasp or seize by wrapping around [1]. Prehension has evolved many times in both animals and plants to enable an organism to grasp vegetation. Examples include elephant trunks, giraffe tongues, plant tendrils, monkey tails, and the human hand. While prehensile behavior has long been noted in these organisms, few systematic studies exist because the action is fast and often performed in treetops. In comparison, elephants are terrestrial, highly trainable, and have an enormous range of weights they can lift. Thus, elephants are ideal for studying the mechanics of prehensile grip.

One of the earliest studies of elephant trunk biomechanics was a 1991 study of an Asian elephant lifting a kettlebell-style weight. Based on the bending of the trunk during the lift, the authors showed the trunk had an effective modulus of elasticity of 985 kPa[2]. Similar to how fish suck prey into their mouths, elephants can inhale air to grab nearby items such as tortilla chips[3]. They can use their trunk “fingers,” the prehensile tips of their trunk, to pack together a series of items for a single pickup [4]. Although the range of movements appears complex, the trunk’s motion can be simplified to a set of 17 basic motion primitives[5]. These primitives were discovered by 3D tracking two adult male elephants grasping objects of varying shape and mass [5]. In this study, we provide an elephant with identically-sized but variously weighted dumbbells to isolate how the elephant changes its gripping strategy.

We next review work on prehension in monkeys and the human hand in order to obtain some context for elephant prehension. Prehensile tails are thought to have evolved in dense South American forests, where animals often traverse narrow supports and distribute their weight to the surrounding canopy [6, 7]. The most well-known examples of prehensile animals are the Atelinae, a subfamily of monkeys that includes howler and spider monkeys [8]. Both spider and howler monkeys can hang their entire body weight (up to 10 kg) from their tail, a behavior that frees their hands to manipulate fruit [9]. This feat requires specialized anatomy: specifically, substantial musculature, innervation to control the tail, and a particular region in the brain for tail control. Another adaptation that makes these monkeys more prehensile than the capuchins is their friction pad, a hairless and highly sensitive strip of skin on their tail [10]. In contrast, capuchins have a tail that is covered in fur. Due to their quick movement through the dense canopy, there are few measurements of tail prehension[11]. Instead of a smooth friction pad as in capuchins, elephants have had a wrinkled ventral section in their trunk [12]. In this study, we consider how wrinkles may improve the surface area of the grip.

Much of the neuroscience of prehensile gripping was found by studying humans. Many tools in the human-built world, such as knobs, steering wheels, and door handles, were designed to be operated by the human hand. Thus, robot designers have shown a keen interest in mimicking the human hand [13]. The human hand has five fingers, over 25 degrees of freedom, and three grasping motor primitives (transverse, perpendicular, or parallel to the palm). Motor primitives are described as neural mechanisms that assist with coordinated motions. Different postures present varying degrees of force, motion, and sensory information [13]. Like the human hand, the elephant can move with precision and high force due to the African elephants’ abundant facial neural control from over 60,000 facial neurons[14].

Elephant trunks and other soft biological structures have been sources of inspiration for soft robotic manipulators for the past twenty years[15, 16, 17, 18]. A novel way of actuating soft robots is through robotic skins, which use pneumatics to move previously inanimate objects [19]. Robotic skins can be used for both actuation, and sensation [20]. Snakes are also prehensile, and their ability to climb trees relies in part on the dynamics of their ventral scales, which can be angled like venetian blinds to increase friction[21]. Bio-inspired robots with actuated ventral scales can climb inclined surfaces using similar mechanisms to snakes[22]. In this work, we will observe how the elephant trunk can make contact with objects using wrinkles along its trunk

## 2 Materials and Methods

### 2.1 Elephant experiments

We initially attempted to train two female elephants at Zoo Atlanta, but only one elephant was receptive. Experiments are thus reported for one elephant, specifically, a 35-year-old female African Elephant (*Loxodonta africana*) of mass 3360 kg and height 2.6 m. We conducted experiments outdoors, at the edge of the elephant’s enclosure at Zoo Atlanta. The experiments occurred over two-hour periods in the mornings of spring and summer 2018 before Zoo Atlanta opened to the public. The staff at Zoo Atlanta supervised all experiments.

To train the elephant to lift, the zookeepers used a reward system beginning with gesturing the elephant towards the barbell (**Figure 1b**). If the elephant accomplished the correct task of grabbing and lifting the bar, food was rewarded(**Figure 1c-d**). If an incorrect outcome was observed, then the experimental procedure was repeated until the trial was successful. Once an 80% success rate was achieved, we commenced weightlifting experiments (**Figure 2a**). It took 15 attempts and 10 minutes of training to accomplish an 80% success rate.

**Figure 1:**
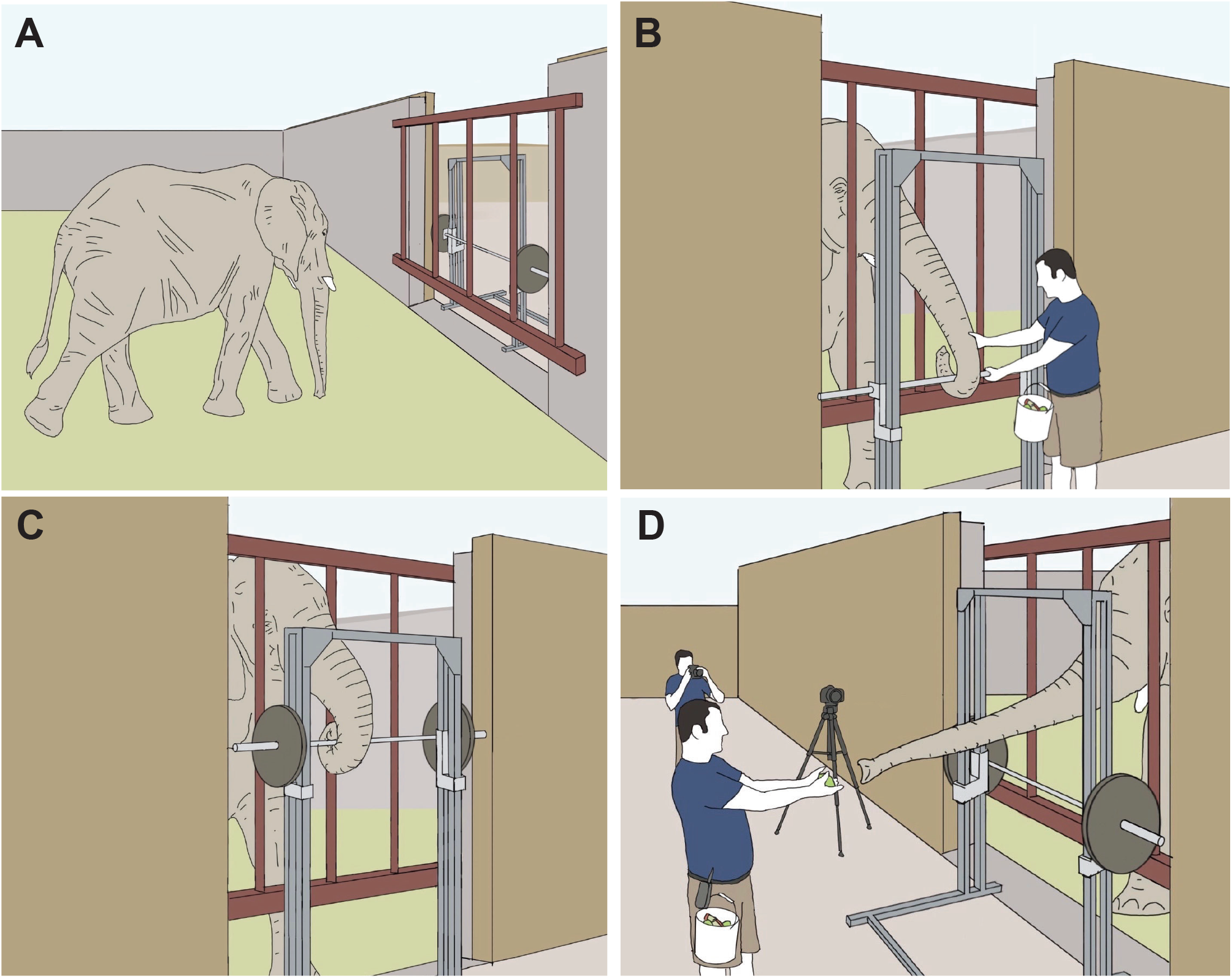
Procedures for elephant lifting the barbell. a) The African elephant *Loxodonta africana* approaches the barbell setup for the experimental procedure. b) A Zoo Atlanta elephant keeper instructs the elephant how to wrap its trunk around the barbell and lift it. c) Elephant completing a trial with a heavier weight. d) After completing a trial, the elephant reaches out to Zoo Atlanta keeper for a food incentive. Illustrations by Benjamin Seleb.

**Figure 2:**
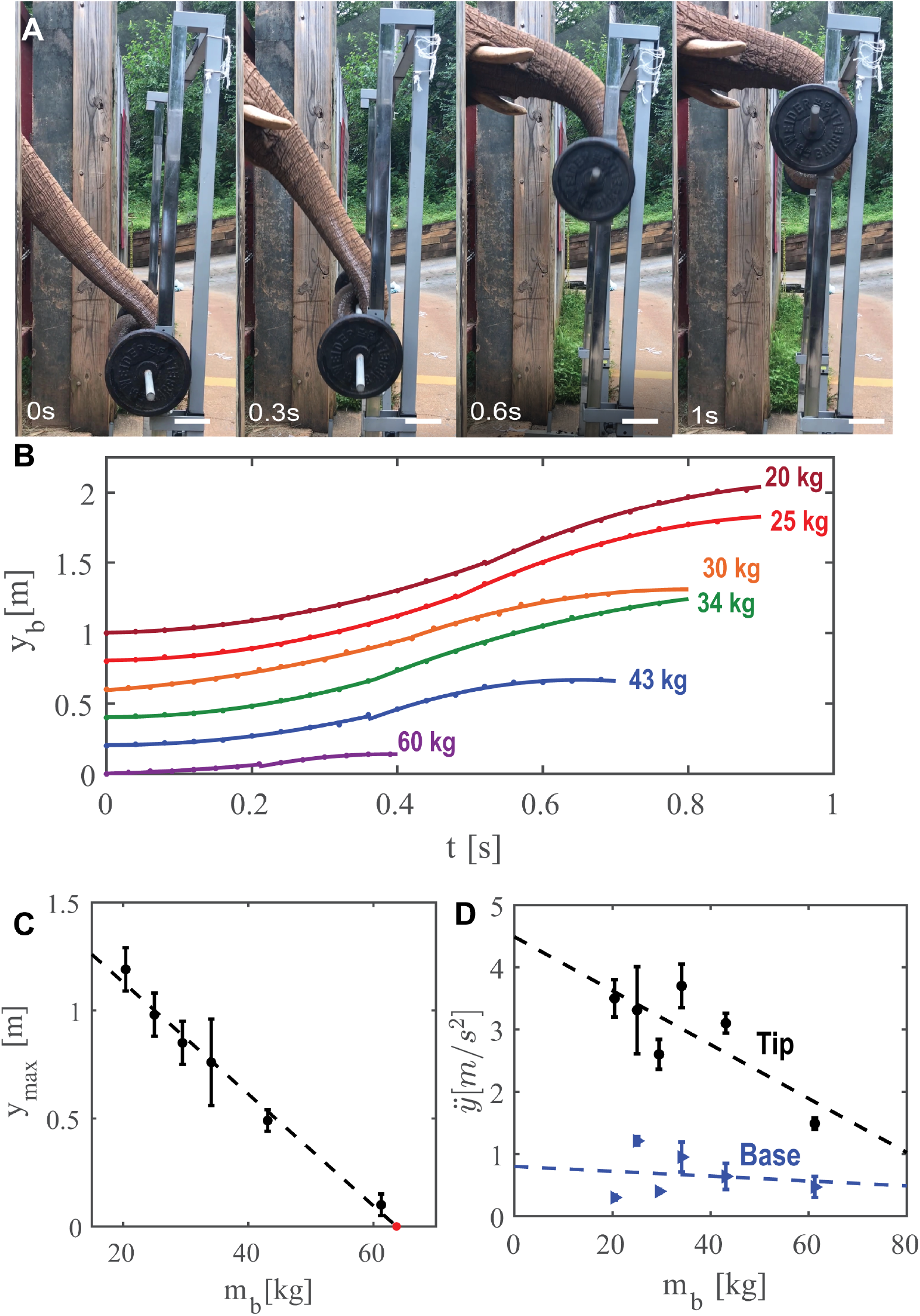
Kinematics of weight lifting. a) Time series of the elephant lifting a barbell at increments of 0.3s with scale bars showing 10 cm. b) Time course of the position of the barbell, with trajectories staggered for clarity. Weights include: maroon (20kg), red (25 kg), orange (30 kg), green (34 kg), blue (43 kg), purple (60 kg). Solid lines are best fit lines associated with constant acceleration *a* and constant deceleration. c) Relationship between maximum height of the elephant trunk and the barbell mass. A red dot indicates the x-intercept, which is the prediction of the maximum mass that the elephant can lift in this setup. d) Relationship between vertical acceleration and barbell mass. The tip and the base of the trunk are shown by black circles and blue triangles, respectively. Best fits given by the dashed lines.

Experiments were conducted with a Smith Machine (Powerline PSM144X, 2.0 × 1.1 × 1.9 m), which uses twin frictionless carriages to constrain the barbell to move vertically. The barbell was placed at a set distance of *w* = 0.5 ± 0.05 m (n=22) away from the enclosure bars. As a result of this distance, the elephant had to rely on its trunk to lift. Without the restraint of the bars, the elephant would likely use a combination of its forehead, trunk, and tusks to lift heavy objects.

Iron weight plates were added to the 20-kg bar to provide the elephant a set of six weights comprising 20, 25, 30, 35, 43, and 60 kg. Note the 43 kg mass results from a 45-pound bar with two 25-pound weights added. The elephant completed four trials of each weight, with each successful trial ending in a food reward and one-minute rest between each lift. When weights were changed, five minutes of rest were given to change the weights and re-secure the frame to the ground using 80 kg of barbell weights.

Twenty-two barbell lifts were filmed using a high-definition digital video camera (Sony HDRXR200) and iPhone 8. We tracked the position of the weight along the 2.0 m height of the Smith machine to accurately determine the barbell height. In two trials, the elephant barely lifted the bar above its original position; such experiments were considered incomplete, and the data were not analyzed. Analysis of the elephant lifting 50 kg was removed from the analysis because the elephant broke the Smith machine during the 50 kg test. During testing, the elephant struggled to lift the 60 kg barbell and only proceeded to lift it twice.

We tracked the trunk tip shape by first drawing along a line of chalk on the mid-line of the right lateral side of the elephant trunk. Using Tracker, an open-source video analysis tool (https://physlets.org/tracker/), we tracked 60 equally spaced points along this line. The speed and acceleration of the elephant lift were determined by tracking the barbell side-view position. Tracking the trunk’s base was impossible because the elephant’s head rose out of the frame.

### 2.2 Dissection of an elephant trunk

Icahn School of Medicine at Mount Sinai, New York provided access to a frozen trunk from a 38-year-old female African elephant, *Loxodonta africana*, that initially lived in a Virginia zoo. Detailed information about the elephant can be found in the pathology report (**Supplemental Figure 1**). The elephant’s body weight before death was approximately 4000 kg, and the weight, age, and sex of the elephant were comparable to those of the elephant filmed in our study. The trunk was cut into several parts and stored in a freezer in 2015 at −15°*C* until we dissected it in July 2016. In January 2018, the specimen’s distal tip was fully thawed and scanned on a Siemens Dual Source Force CT to measure the trunk’s nasal passageways and outer diameter. A helical scan was performed with 80 kV, 183 mAs, a slice thickness of 0.5mm, an acquisition speed of 737 mm/sec, and a temporal resolution of 66 msec (**Figure 3a-b**, **Supplemental Video 4**). We scanned the distal portion of the trunk up to around 110 cm from the tip. That was as much of the trunk as that could fit in the CT scanner. We obtained 27 measurements of the trunk’s inner diameter as the scan progressed from the proximal root to the distal tip (**Figure 4c**). We also rendered the entire CT image of the trunk to see the three-dimensional structure (**Supplemental Video 5**).

**Figure 3:**
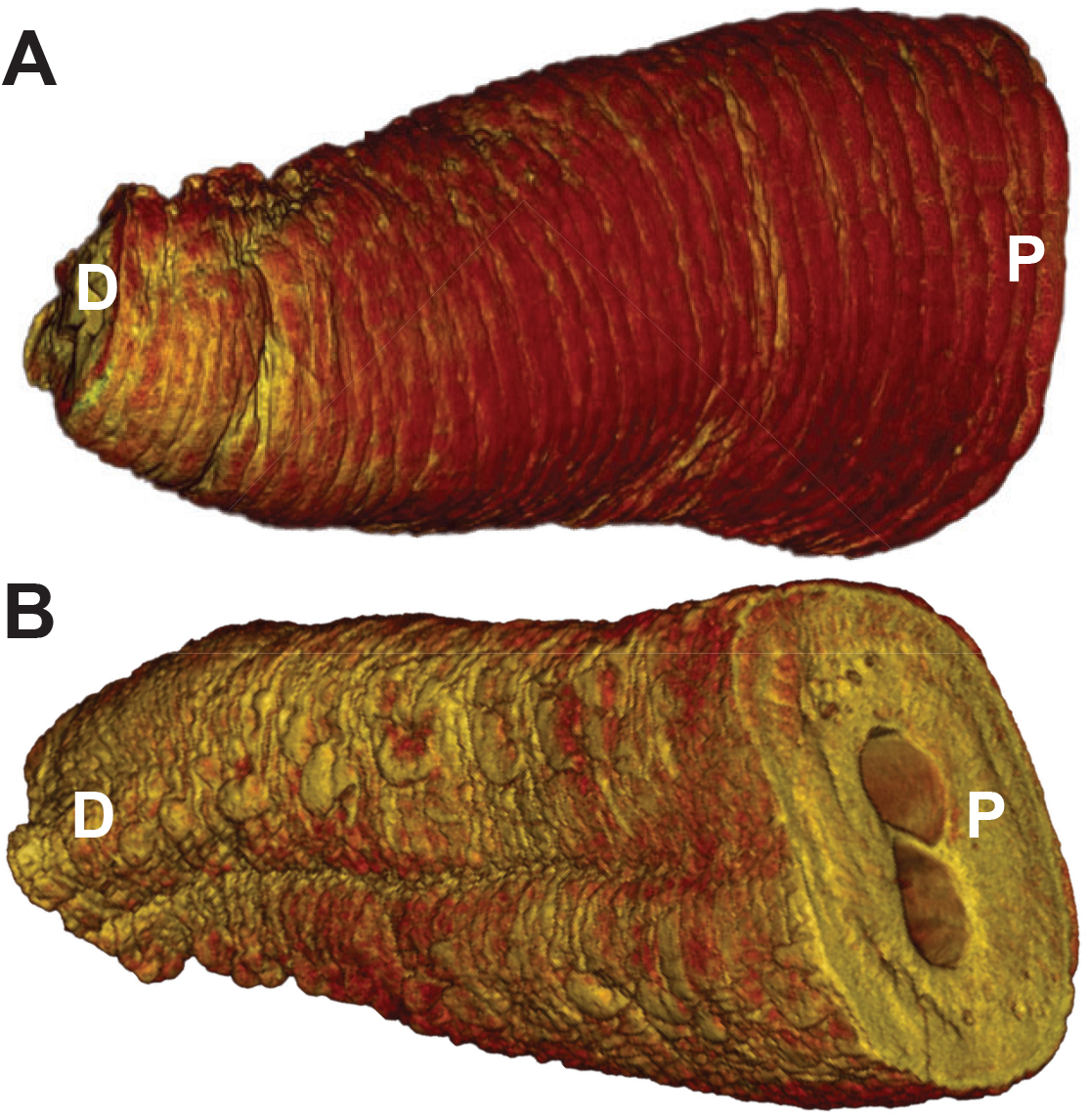
CT scan of the trunk of a 38 year old female African elephant *Loxodonta africana*. a) dorsal section and b) ventral section. P refers to proximal (towards the skull), D refers to distal (towards the tip). The section shown is the distal 60-cm of the trunk.

**Figure 4:**
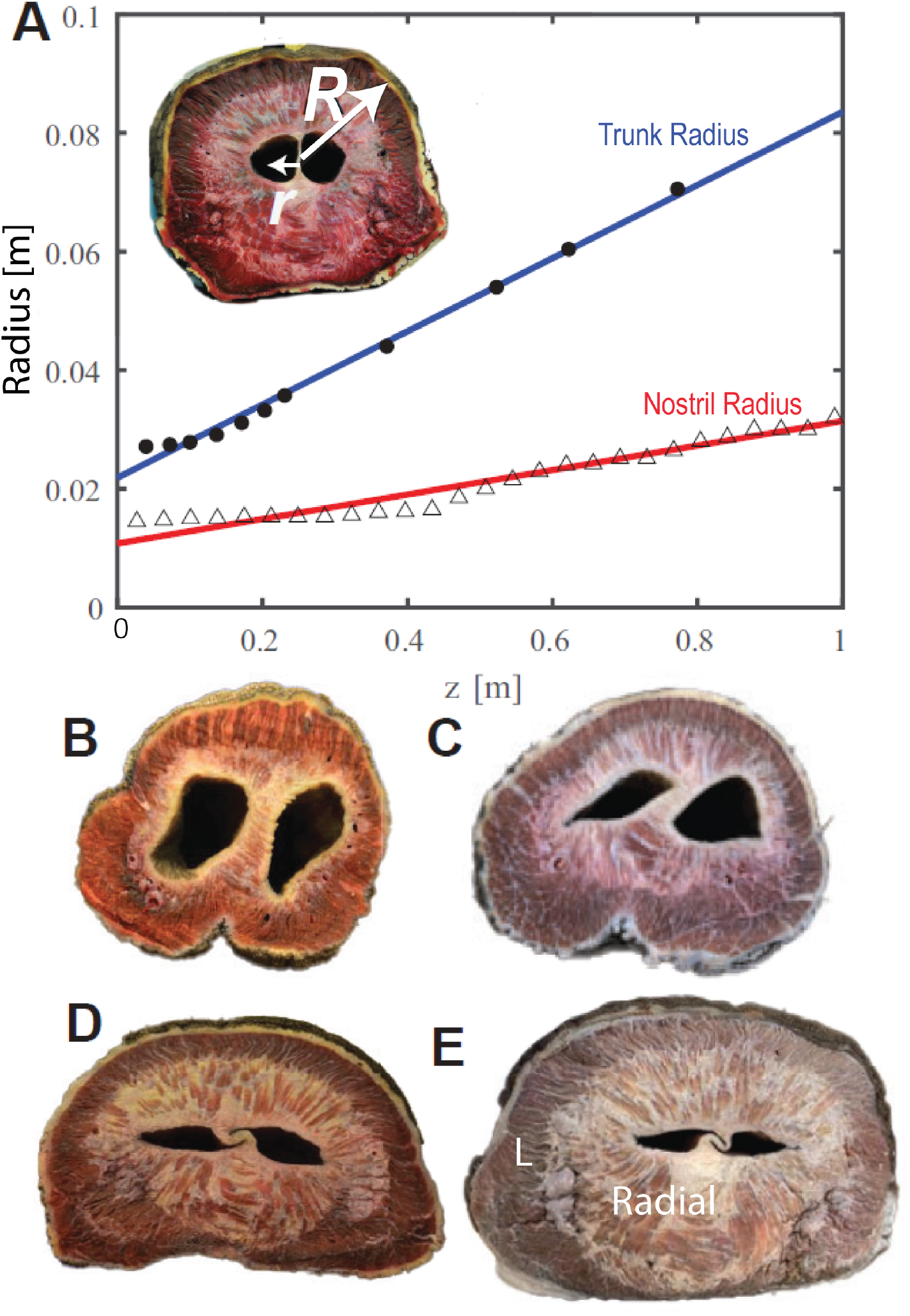
Elephant trunk anatomy. a) The relationship between radii of the trunk and distance *z* from the tip. The trunk outer radius, *R* is given by the shaded circles and inner nasal radius, *r* by open triangles. Linear best fits are shown by the solid lines. b-e) Elephant trunk cross sections displaying muscle fibers and negative space created by nasal passageway. b) cross section 28 cm from distal tip. c) 56 cm from distal tip. d) 100 cm from distal tip. e) 140 cm from distal tip With “L” indicating longitudinal muscles, and “radial” indicating radial muscles.

## 3 Mathematical Modeling

### 3.1 Elephant trunk geometry

We modeled both the elephant trunk and its nasal cavities as conical frustums [23]. Assuming the mass density of the trunk is *ρ* = 1180 kg/*m*^3^, measured from a cross-section of an elephant carcass’ trunk [2], the mass *m*_*t*_ of a trunk segment of length *z* may be modeled using a solid frustum with two hollow frustums as nasal cavities. With these assumptions, the mass may be written [24]:

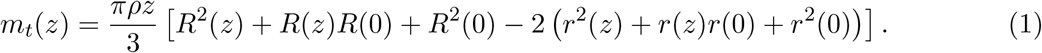

The length *z* is measured from the trunk tip, and *R*(*z*) and *r*(*z*) are, respectively, the outer and inner radii of the trunk at a position *z*. Based on the frozen trunk measurements, the inner and outer radii for the trunk tip are *r*(0) = 1.1 cm and *R*(0) = 2.2 cm, respectively. The nasal passages of the frozen trunk were squashed by self-weight. We used the nasal circumference to extrapolate the inner radius (**Figure 4d**).

### 3.2 Tension and power applied

To determine the force required to lift the barbell, we consider a vertical force balance on the trunk tip. A control volume is shown schematically in **Figure 5a**. When the barbell is lifted, the elephant lifts both the barbell and the trunk itself. The total mass to be lifted is *m* = *m*_*t*_ + *m*_*b*_ where *m*_*b*_ is the barbell mass and *m*_*t*_ is the mass of the trunk segment in contact with the barbell. In reality, the elephant’s proximal portion of the trunk rises as well. However, the average height of the trunk segments not in contact cannot be calculated since part of the elephant trunk leaves the video frame. Thus, a weakness of our method is that the applied force and power will be underestimated. To determine the vertical acceleration at the base of the trunk, we measured the vertical position of the apex of the tusk, which stayed within the video frame.

**Figure 5:**
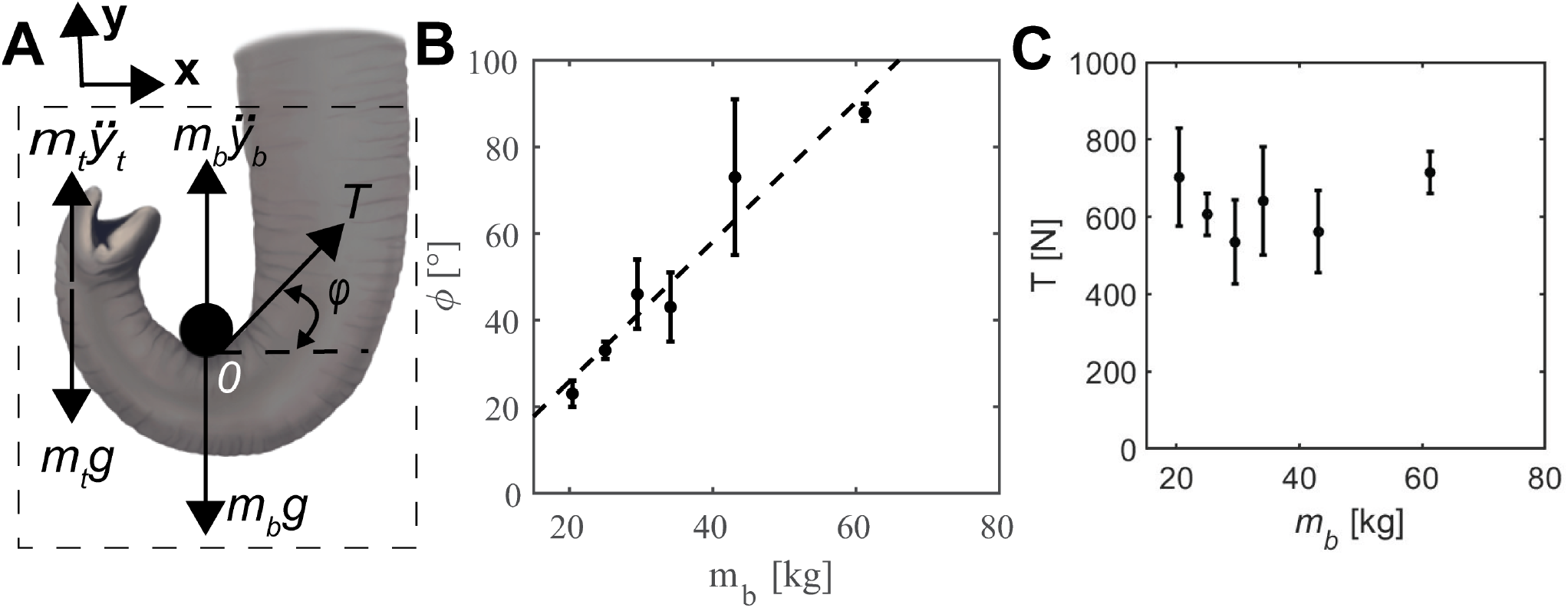
Forces exerted on the barbell. a) Free body diagram of the elephant lifting a barbell. The elephant applies a tension *T* to the barbell to lift it at contact point *O*. The trunk is at an angle of contact *ϕ* with respect to the horizontal. The combined trunk and barbell mass experiences gravity *g* and an upward acceleration 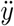. b) The relationship between the angle of contact *ϕ* and barbell mass. Linear best fit is given by the dashed line. c) The relationship between the calculated tension *T* and barbell mass.

The trunk exerts a tension *T* to lift. The angle *ϕ* between the trunk and the horizontal is measured at the instant the lift begins (**Figure 5a**). We neglect the displacement in the horizontal direction because the Smith Machine constrains the barbell from moving vertically. Since the trunk segment is wrapped around the barbell, both move with the same vertical speed 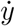 and acceleration 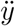. By Newton’s law, the vertical force balance may be written

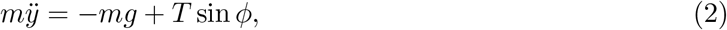

where *g* is the acceleration of gravity and *T* is the tension applied. Solving Equation (2) with respect to the tension force *T* yields

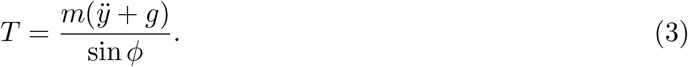

Thus, by measuring the angle *ϕ* and the acceleration 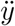, we can estimate the force exerted to lift the barbell.

We calculate the power to lift two parts of the trunk, the tip, and the base, which is assumed to be the apices of the tusks. Each of these parts has its own mass that is estimated from Equation (1). The average power exerted to lift part *i* of the trunk may be written as the ratio of the gravitational potential energy and duration *t* of lift. The gravitational potential energy of a trunk segment of mass *m*_*i*_ is written as *U* = *m*_*i*_*gy*_*i*_, where *y*_*i*_ is the change in the height of the center of mass of that portion of the trunk. The mass of the tip was assumed to be between 5.4 and 9 kg, depending on the observation of the length of wrap. The mass of the trunk base was assumed to be 60 kg, which we give as an upper bound. The power is thuswhich is consistent with the definition of power for human weight lifters [25].

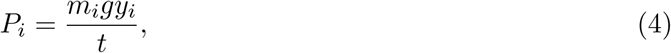

### 3.3 Contact area

While the dorsal trunk is cylindrical, the ventral part of the trunk is planar and covered with wrinkles. To estimate the contact area of the trunk and the barbell, we measure the frequency *ω* and amplitude *A* of the trunk wrinkles at 80 positions, including eight axial positions along the frozen elephant trunk and 10 azimuthal positions for each axial position. Previous work showed that the trunk tip is nearly inextensible: the distal 30 cm of the trunk stretches less than 10% strain, which is small compared to the stretch mid-distally, which is 25 percent[12]. We thus assume that the wrinkle geometry of the frozen trunk matches the live elephant. Assuming a sinusoidal wrinkle profile, the radius *R* of the ventral trunk skin as a function of distance *z* from the tip is written in the results section in Equation (4.3).

We report the amplitude *A*(*z*) and frequency *ω*(*z*) in the results section and in **Figure 7E-F**. To calculate the surface area of contact of a trunk segment, we utilize the arclength formula which states the arclength *s* of the trunk segment is

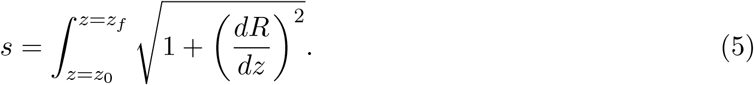

This integral is calculated numerically using MATLAB ode45, between the two points *z*_0_ and *z*_*f*_, which defines the trunk segment in contact with the bar. We assume the contact region is a wrinkled planar trapezoid of height *s*, and lengths *D*(*z*_*f*_) and *D*(*z*_0_), which are the diameters of the trunk at the points *z*_*f*_ and *z*_0_. The area of this trapezoid is :

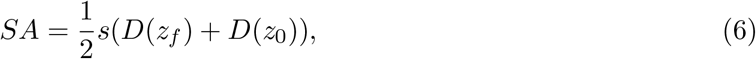

where we have assumed as an upper bound that the entire wrinkle surface area contacts the bar.

## 4 Results

### 4.1 Trunk geometry

To calculate the force applied by the trunk, we first measure its shape. We characterize the frozen trunk from the Smithsonian using CT-scanning and dissection (**Figure 3, Supplemental Video 4**). At the proximal base, the cross-section is dominated by radial muscle, marked “radial,” the light-colored muscle close to the nasal cavities. A large amount of radial muscle is presumably to help with lifting as the base does not stretch much longitudinally (**Figure 4e**) [12]. The longitudinal muscle marked “L,” is darker red and lies between the radial muscle and the skin of the trunk. The proportion of radial muscle shrinks progressively towards the distal tip, while the proportion occupied by the nostrils increases. The distal tip of the trunk lacks radial muscle and is instead dominated by two oblique muscle groups (**Figure 4b**), which assist with wrapping around objects. The trunk is a hollow conical frustum permeated by a pair of nostrils. From the CT scans, we obtain a relationship for both the inner and outer radius of the trunk at a distance from the tip, *z* (**Figure 4a**). The inner radius *r* is given by the open triangles and the outer radius *R* by the closed points. The solid lines are linear least squares best fits given by

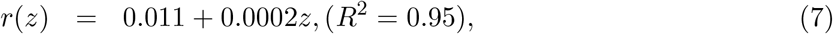

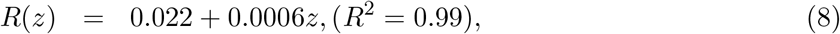

with all units in meters. At the tip, the inner and outer radii are 1.1 cm and 2.2 cm. At a point 100 cm from the tip, the trunk has inner and outer radii of 3 cm and 8 cm. Using Equation (1), we calculate a trunk mass of 97 kg, which is close to the experimental measurement of 110 kg. The mass *m*_*t*_ of the trunk segment in contact with the barbell was calculated for each experiment based on the length in contact. The mass of the trunk in contact ranged from 5.4 kg for the lightest barbell up to 9.0 kg for the heaviest.

### 4.2 Lifting force

The elephant lifts the barbell by first wrapping its trunk tightly around it (**Figure 1**). The trunk arches like a bending beam as it lifts (**Figure 2a** and **Supplemental Video 1-3**). The total energy expended for each barbell weight is related to the maximum height of the lift *y*_*max*_, shown in **Figure 2c**. The linear best fit, shown by the dashed line, is

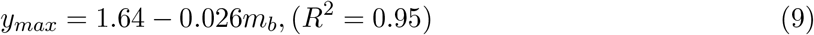

where *y*_*max*_ is the height lifted in meters. For this and future equations, *m*_*b*_ is the weight of the barbell in kg. The elephant lifted the lightest weight to a height of 1.19 ± 0.1 m (n=4), nearly touching the top of the weight rack, and the heaviest weight to less than one-tenth the height at 0.1 ± 0.05 m (n=2). Clearly, the elephant lifted heavier weights less. The x-intercept of Equation (9) shows that that the heaviest weight that elephants can lift in this setup is 63 kg, which is just 2% of its body weight and 65% of its trunk weight. This weight is less than we expected given an elephant’s feats of strength. We surmise that the barbell apparatus constrained the trunk motion to the vertical, preventing the elephant from using body weight to push or lift the object.

The time course of the barbell height *y*_*b*_ is shown in **Figure 2b**, where we spaced out the trajectories for clarity. After the trunk wrapped around the barbell, the lifts were fast, taking approximately 0.5 − 0.8 s across the weight classes. Each function was fit with two quadratic best fits separated by an inflection point between the acceleration and deceleration phases. The inflection point usually occurs at the midpoint of the lift. In the acceleration phase, the position may be written as *y*_*b*_ = *at*^2^ where *t* is time, and *a* is the acceleration. This equation describes lifting from a rest position with a constant acceleration *a*. Such a relationship fits the acceleration phase well, with an *R*^2^ greater than 0.95. The deceleration phase was fit with the position and velocity at the inflection point, as well as a constant deceleration: *y*_*b*_ = *y*_0_ + *vt* − *bt*^2^. We found no clear trend between deceleration and barbell mass, so the deceleration was not reported. The fits for the entire lift are shown in **Figure 2b**. The acceleration 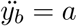 for each barbell mass is shown in **Figure 2d**, with the linear best fit given by

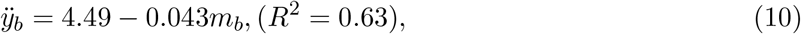

where 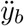 is in m/s^2^ and *m*_*b*_ is in kg. Equation (10) indicates that an elephant has a lower acceleration for heavier weights: acceleration ranges from 3.9 m/s^2^ for the lowest weight to less than half that value for a weight three times heavier.

Closer to the elephant’s head, lifting is accomplished by some combination of rotation and lifting of the head. The red points in **Figure 2d** shows the vertical acceleration of the trunk base: the acceleration of 1 m/s^2^ is small compared to most of the trunk tip accelerations. We conclude that vertical motion is small, and instead, the neck acts like a fulcrum to provide rotational motion to assist lifting by the trunk tip.

To calculate the tension applied by the trunk, we measured the angle that the trunk intersects the barbell. **Figure 5b** gives the angle *ϕ* of the trunk with respect to the horizontal, where the dashed line is the least squares linear best fit. The relationship between the angle of the trunk and barbell mass is

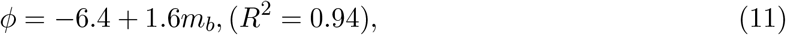

with *ϕ* in degrees. An angle of 90 degrees indicates that the elephant orients its trunk vertically. The elephant increases the angle of contact *ϕ* from 23 ± 3° (n=4) for the lightest weight to nearly four times that amount, or 89 ± 2° (n=2), for the heaviest.

Given the angle *ϕ* and the length of the trunk *z* wrapped around the bar, we calculate the trunk tension *T* using Equation (3) in the Math Methods. The relationship between tension and the mass lifted is shown in **Figure 5c**. Although the barbell weights increase by a factor of three, the tension increases by only 20%, maintaining an average value of 620 ± 64 N (n=22) across all trials. We thus see that elephants have dual “strategies” to maintain tension when lifting heavy weights: they decrease acceleration and orient their trunk more vertically. These strategies may not be volitional: they may simply arise from trying to lift a heavier weight with a finite muscle of limited force and power. Nevertheless, we see the trunk adapts to different postures and kinematics for different weights.

**Figure 6b** shows the relationship between power exerted and the mass lifted, with red points referring to the trunk base and black points to the trunk tip. These powers are overestimated because they only consider the maximum deflection of the highest point rather than the center of mass of each section. The power expenditure of the tip is U-shaped, with a peak power of 357 ± 79 W (n=4) for intermediate masses. No matter what weight is lifted, the power expended at the trunk base remains higher than the trunk tip. This is because the mass of the base is 60 kg whereas the mass of the tip is less than 10 kg.

**Figure 6:**
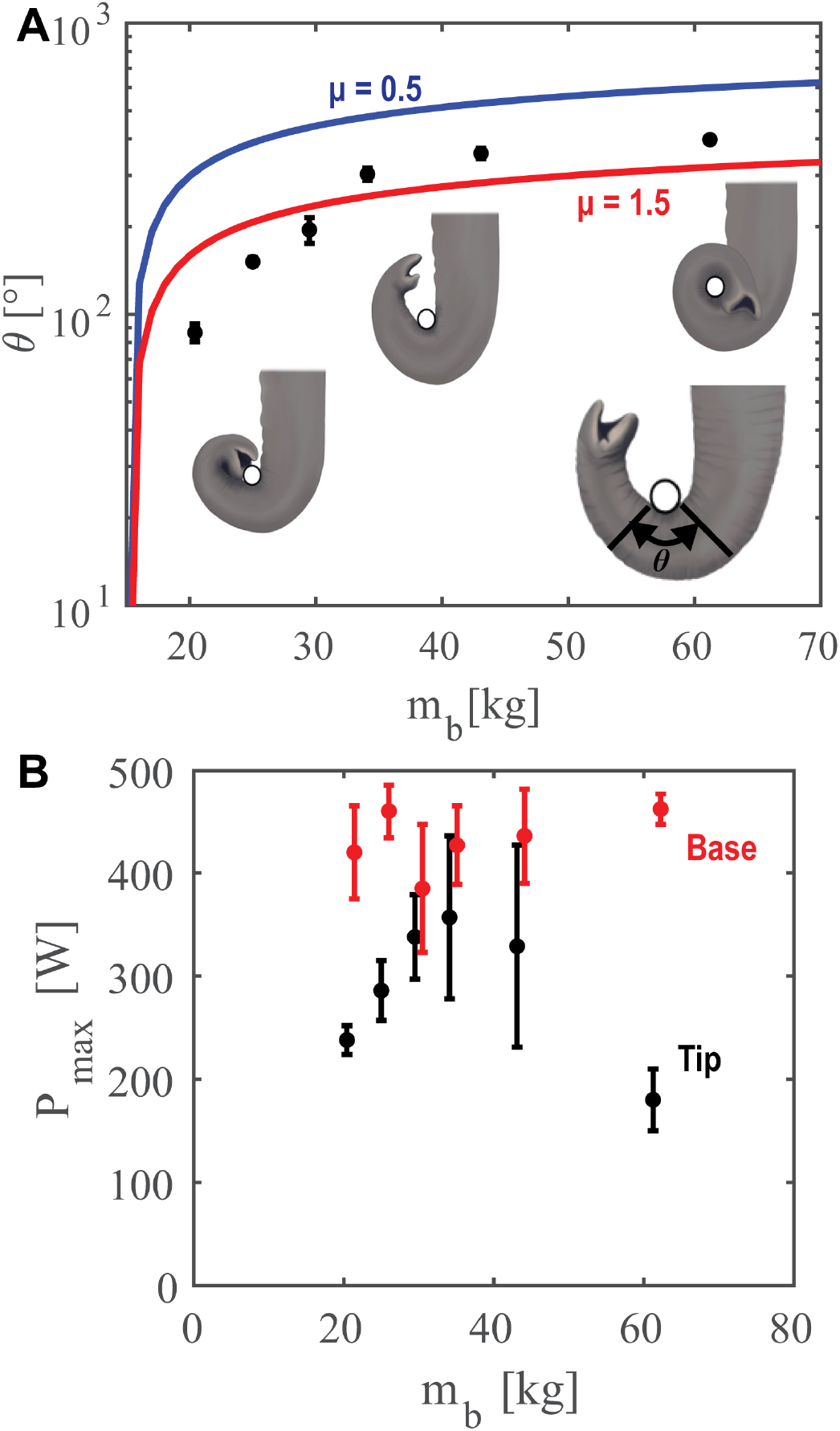
Elephants wrap the trunk to lift heavier weights. a) The relationship between angle of trunk wrap *θ* and barbell mass. Schematics, from left to right, show the increasing wrap of the trunk for barbell weights 20 kg, 25 kg, and 60 kg. Theoretical predictions with friction coefficients of 0.5 and 1.5 are shown by the blue and red lines, respectively. b) Maximum power exerted to lift different barbell masses. Power is calculated at two locations, the distal tip (black circles) and the proximal root (red circles).

### 4.3 Prehension

Although the elephant was at a constant distance to the bar, it wrapped a greater length of its trunk to lift heavier weights. **Figure 6a** shows the progression of trunk wrapping, from *θ* = 87 ± 6° (n=4) to 400 ± 12°(n=2), an increase in wrapping angle of 400%. To lift the lightest weights (*m*_*b*_ = 20 and 25 kg), the trunk’s distal tip extended past the barbell and wrapped around the bottom half, creating a lip that kept the barbell in place. When lifting medium weights (*m*_*b*_ = 30 − 43 kg), the trunk extends further, using a thicker section of its trunk to wrap. Finally, when the elephant lifted the heaviest weight (*m*_*b*_ = 60 kg), the trunk wrapped 413 degrees or more than a full cycle. Wrapping a greater angle increases the contact area between the barbell and the elephant trunk. We note that the largest angle supporting the barbell’s weight is 180 degrees, corresponding to the bottom half of the barbell. Any additional wrapping helps with stability rather than weight support.

For us to rationalize the increased wrapping angle with heavier weights, we consider the capstan, a rotating device that amplifies a sailor’s ability to pull a rope [26]. The classical capstan model shows that *T*_*b*_*/T*_*a*_ = *e*^−*μθ*^ where *θ* is the angle subtended by the capstan, *μ* is the coefficient of friction, and *T*_*b*_*/T*_*a*_ is the ratio of the sailor’s force to the force on the other end of the rope. Assisted by the friction on the rope wrapped around the capstan, the sailor can amplify its force *T*_*b*_ to support a load *T*_*a*_.

Applying the capstan problem to the barbell wrapping, we may consider the “sailor” to be *T*_*b*_ = *m*_*t*_*g*, which is the gravitational force imposed by the trunk pendent mass *m*_*t*_ wrapped around the barbell. The weight of the pendent, as well as the friction at the contact area, opposes the barbell weight *T*_*a*_ = *m*_*b*_*g*. By wrapping greater angles, the elephant can use the weight of the pendant to avoid losing grip on the barbell as it is lifted. Based on the arclength of the trunk wrapped, the weight of the pendent mass varies from 5.4 to 9 kg and increases with barbell mass. Simplifying the capstan model, we find

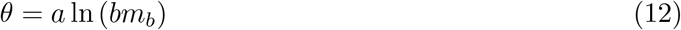

where 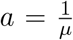 and 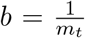. A least-square best fit, given in red, fits the data quite well, showing that 100 − 300 degrees of wrap is sufficient to hold the barbell (**Figure 6a**). The free parameter for the best fit is a high but still physically reasonable friction coefficient of *μ* = 1.5, comparable to the friction coefficient of 1.6 for bio-mimetic snake robots scales on styrofoam [22]. Snakes can change the angle of their ventral scales to increase frictional forces as they climb tree trunks and other vertical surfaces. Such actively deformable surfaces are analogous to the friction-enhancing wrinkles and coarse hair on the trunk. For comparison, we show in blue another wrapping angle using the friction coefficient of human skin on metal (*μ* = 0.8). Indeed, such a low friction coefficient requires 200-500 degrees wrapping angles, which are higher than the observed angles. Both models bound the data and give evidence that the combination of self-weight of the pendant mass and skin friction prevent the barbell from slipping as it is lifted.

Our capstan model assumed a constant friction coefficient, but the trunk may be able to modify its friction coefficient using its wrinkled grip, as shown in **Figure 7A**. We measured by hand the wrinkle amplitude *A* and wavelength *λ* as a function of the distance *z* from the tip. A linear least squares best fit shows that wrinkles increase both their amplitude and wavelength with distance from the tip:

**Figure 7:**
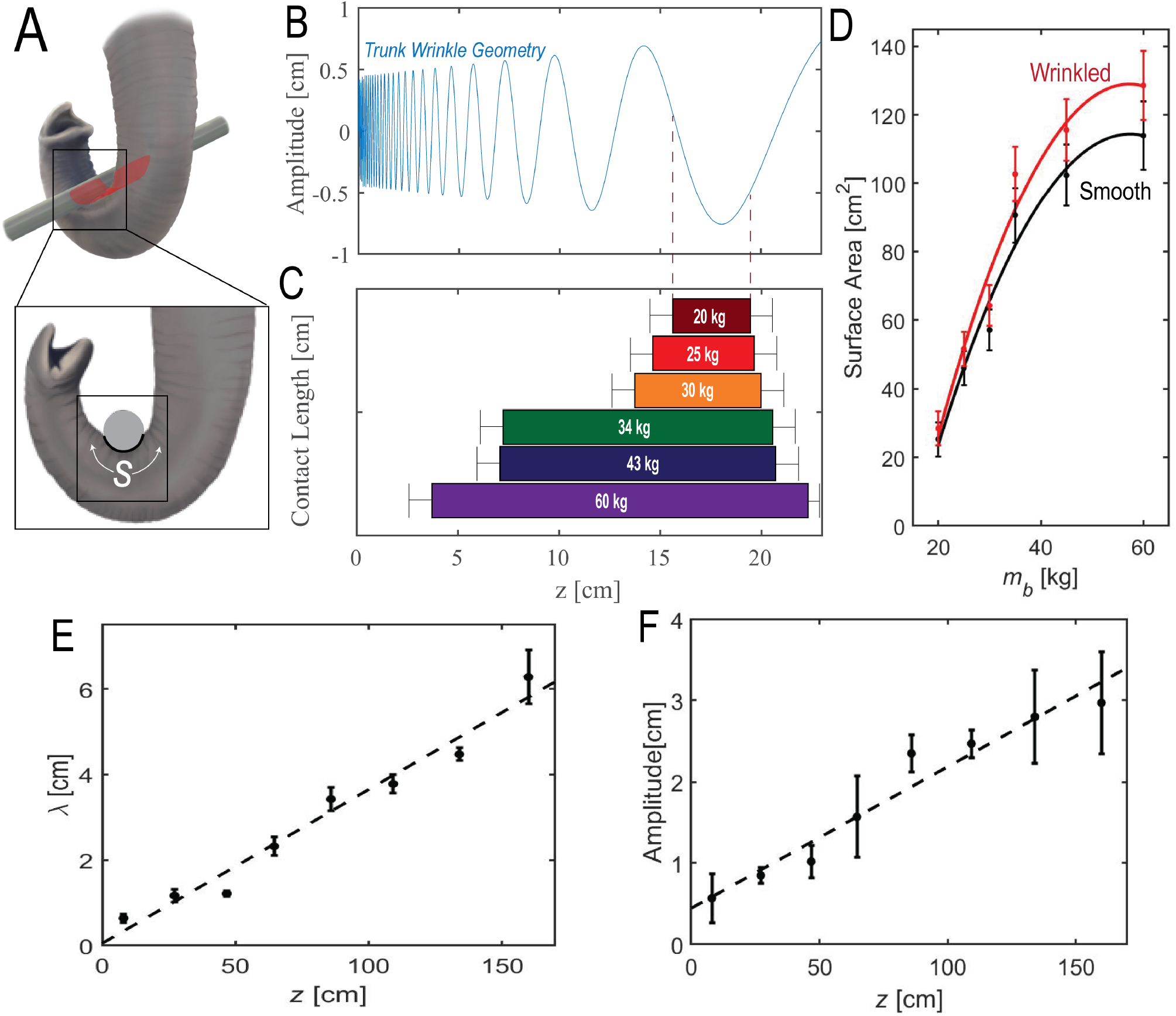
Elephant wrinkles increase area of contact with the barbell. a) Schematic displaying the elephant’s area of contact with the barbell using its wrinkled ventral trunk. b) Ventral surface profile along the trunk’s long axis. Wrinkles increase in amplitude and wavelength with distance from the tip, which is at *z*=0. c) Observed contact length *s* of the barbell for different barbell weights. This length does not take into account wrinkles. d) Surface area of contact across weight classes, with black showing the surface area without wrinkles, and red the surface area with wrinkles. e) Wavelength of the elephant wrinkles from the tip of the trunk to the base. f) Amplitude of the elephant wrinkles from the tip of the trunk to the base.

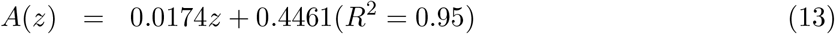

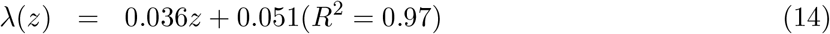

(**Figure 7E-F**) where *A, λ*, and *z* are in cm. Assuming that the trunk surface has a sinusoidal wrinkle pattern, the wrinkle height may be written

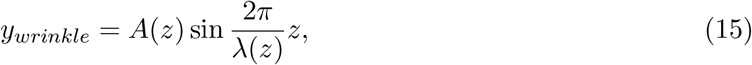

and is shown in **Figure 7B**. Using these relationships, we use equation Equation (6) to estimate the surface area of the wrinkled skin. As the barbell mass increases, the elephant wraps with increased arc length and greater surface area (**Figure 7C**). We consider two contributions to its surface area. First, we consider a smooth ventral trunk devoid of wrinkles, shown by the black points (**Figure 7D**).An upper bound for the increase in surface area is that the entire wrinkle, including the peaks and troughs, are wrapped around the bar: The wrinkled surface area, from Equation (6) is shown in red. In reality, the peaks will be compressed, and there will be air gaps in the troughs of the wrinkles. However, further analysis is not feasible without measuring the contact area precisely. The true contact area will like between the black and red curves. When wrapping is low, since the trunk tip has small wrinkles, its additional surface area is low. However, for the heaviest weight, the wrinkles may contribute up to 15% of the surface area (**Figure 7D**).

## 5 Discussion

Clearly, the Smith machine we used was designed for human weight lifting, but it worked for elephants because the power generated by the elephant trunk is comparable to human power. When humans lift a barbell for a power clean, which involves lifting a barbell from the ground to the shoulders, they can achieve a power of 900 W on free weights, and 770 W on machine cleansa for lifting just a 20 kg weight [25]. When using just its trunk, the elephant lifts 20 kg using only 238 ± 14 W of power, but could probably increase this amount with training.

To lift heavier weights, the elephant recruits a greater surface area of contact using its trunk wrinkles. The ridges on human fingertips have been shown to increase friction by two mechanisms [27]. On rough surfaces, the ridges deform and interlock into an uneven surface when gripping surfaces. Our videography was not close enough to observe any deformation of the wrinkles. However, the mechanism seems plausible since elephants often pick up rough objects such as tree bark, whose asperities on the mm to cm seem comparable to those of elephant wrinkles. The other mechanism for human fingertips is more subtle, involving the maintenance of an optimal layer of sweat between the ridges. The length scale of elephant wrinkles is much larger than human fingertip ridges; moreover, elephants have very few sweat glands [28]. Therefore it is unlikely that moisture plays a role in elephant grip.

The use of wrinkles to increase contact area may be useful for improving the grip of soft robots. Artificial muscles composed of silicon elastomers can actuate rigid grippers to pick up objects. One artificial muscle, known as hydraulically amplified self-healing electrostatic (HASEL)[15], can grasp various objects and use its own feedback to estimate the object’s size [29]. Fluid-filled toroidal tubes can grip, catch, and convey objects [30]. Generally, such robots are covered in smooth skin. The addition of wrinkles[12] may improve grip without increasing the squeezing force and risk damaging fragile objects. The wrinkles may also improve the devices’ reach by enabling the skin to stretch. Uncontrolled grippers that rely on the entanglement of filaments could also be covered in wrinkles to improve grip[31]. One day, wrinkled skin could also improve the ability of soft robots to perform sensing. Adding smooth artificial skin to earthworm-inspired robots facilitated feedback control[20]. The addition of wrinkles may enable a more stretchable interface with its subterranean environment.

Many aspects of lifting remain poorly understood. In many prehensile animals, the surface of the skin is heavily innervated with sensors. Elephant skin is substantially tougher than other animals but somehow maintains a sensitive touch. Since our experiments were performed with only one elephant of one sex, further work is needed to generalize our observations across elephant species. African and Asian elephants differ in their trunk anatomy in that African elephants are browsers, and Asian elephants are grazers. This specialization gives African elephants two prehensile trunk fingers, whereas Asian elephants have one, which leads to differences in facial motor control neurons[14].

## 6 Conclusion

In this study, we elucidate the kinematic and gripping strategies of an elephant lifting barbells. As the barbell increased in weight, the elephant maintained nearly constant tensile force by orienting its trunk vertically and accelerating less. The elephant wrapped its trunk around the bar more for heavier weights, presumably to stabilize its grip. We showed that the self-weight of the trunk might be used like a sailor’s capstan to prevent slipping of the barbell. Incorporating a greater length of the trunk also brings into play deeper and longer-amplitude wrinkles, which we believe increase friction. Since one elephant was studied, it remains unknown whether these results generalize to other lifted objects or other elephants. We hope that this work inspires new kinds of adaptable biologically-inspired grippers.

## Data Access Statement

We have included MATLAB files for the elephant lifts, raw elephant lifting trials, and data spreadsheets of strain tracking available for download on a GitHub repository that is linked in the supplemental documents.

## Ethics Statement

In working with animals from Zoo Atlanta. We received research permits from both Zoo Atlanta, permit name, *Weight lifting experiments by elephants*, and Georgia Institute of Technology IACUC, permit number A16032.

## Acknowledgements

This work was supported by the US Army Research Laboratory and US Army Research Office Mechanical Sciences Division Complex Dynamics and Systems Program, under contact number W911NF-12-4-00111. We thank A. Lee, M. Chan, and Y. Zhang for their early contributions. We thank the Zoo Atlanta elephant keepers with their assistance in performing experiments. We thank Dr. Ali Nabaviziadeh for arranging the collaboration with Dr. Reidenberg. We thank the imaging time donated by Dr. Cheuk Tang’s group, Radiology, Neuroscience, & Psychiatry Translation and Molecular Imaging Institute at Icahn School of Medicine at Mount Sinah. We thank J. Ososky and the Smithsonian Institution Museum of Natural History for their assistance with information regarding the frozen elephant trunk as well as loaning the elephant trunk to Icahn School of Medicine at Mount Sinah.

