## Supplemental Tables & Videos for "Elephant trunks use an adaptable prehensile grip": SchulzSubmissionSupplement.pdf

### 1 Supplemental Material

| $m_b$ [kg] | n | $y_{max}$ [m] | $P_{base}$ [W] | $P_{tip}$ [W] | $\theta$ [°] | $\phi$ [°] | T [kN] |
|---|---|---|---|---|---|---|---|
| 45 | 4 | $1.19 \pm 0.09$ | $420 \pm 45$ | $238 \pm 14$ | $87 \pm 6$ | $23 \pm 3$ | $703 \pm 127$ |
| 55 | 4 | $0.98 \pm 0.055$ | $385 \pm 26$ | $286 \pm 29$ | $150 \pm 5$ | $33 \pm 2$ | $608 \pm 54$ |
| 65 | 4 | $0.85 \pm 0.11$ | $427 \pm 62$ | $338 \pm 41$ | $195 \pm 20$ | $46 \pm 8$ | $536 \pm 108$ |
| 75 | 4 | $0.76 \pm 0.19$ | $427 \pm 38$ | $357 \pm 79$ | $303 \pm 15$ | $43 \pm 8$ | $643 \pm 140$ |
| 95 | 4 | $0.49 \pm 0.043$ | $436 \pm 46$ | $329 \pm 98$ | $357 \pm 15$ | $73 \pm 18$ | $564 \pm 106$ |
| 135 | 2 | $0.095 \pm 0.045$ | $462 \pm 15$ | $180 \pm 30$ | $400 \pm 11$ | $88 \pm 2$ | $721 \pm 55$ |

**Table 1:** Supplementary Table displaying the results as expressed in the paper with each column representing the data for each of the weight classes as described by $m_b$ with the columns displaying mean$\pm$ standard deviation for the sample size of $n$

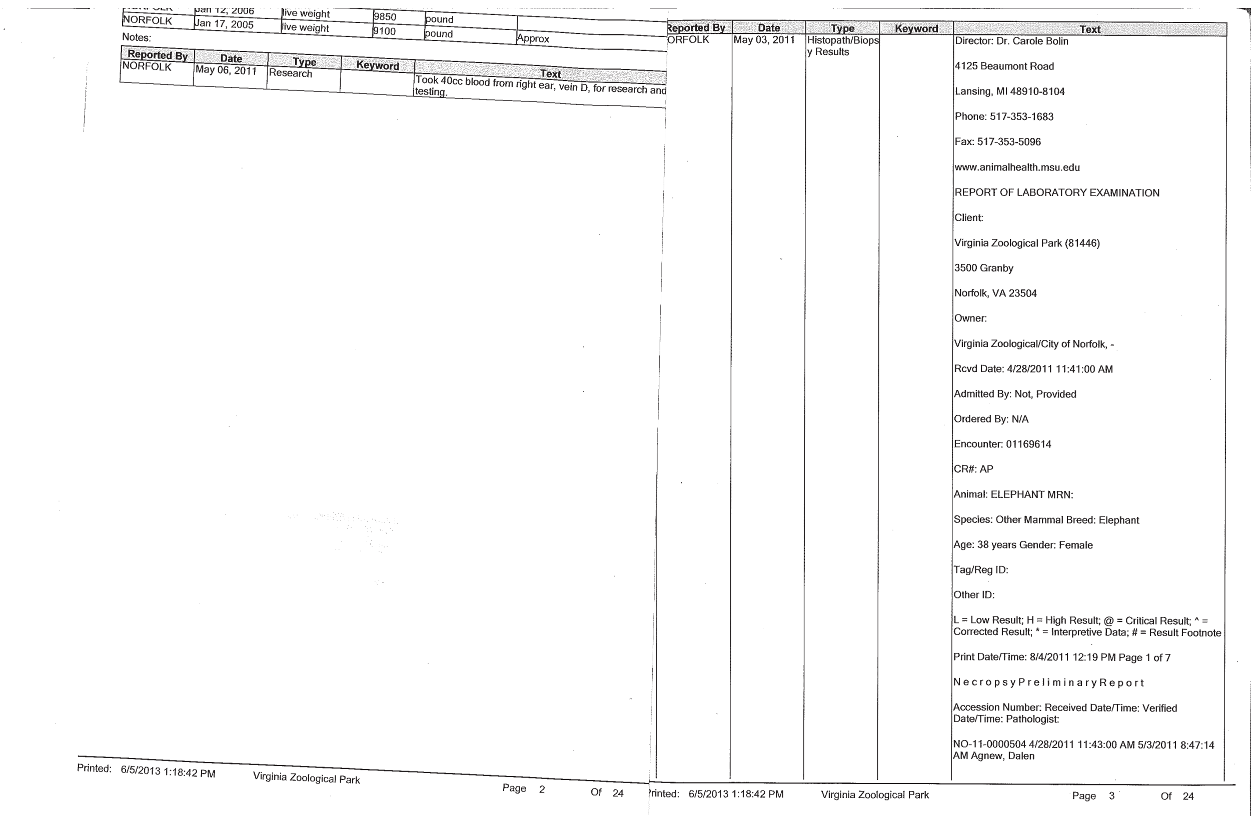

| | | | | | |
|---|---|---|---|---|---|
| [illegible] | Jan 12, 2006 | live weight | 9850 | pound | |
| NORFOLK | Jan 17, 2005 | live weight | 9100 | pound | Approx |

Notes:

| Reported By | Date | Type | Keyword | Text |
|---|---|---|---|---|
| NORFOLK | May 06, 2011 | Research | | Took 40cc blood from right ear, vein D, for research and testing. |

Printed: 6/5/2013 1:18:42 PM Virginia Zoological Park Page 2 Of 24

| Reported By | Date | Type | Keyword | Text |
|---|---|---|---|---|
| NORFOLK | May 03, 2011 | Histopath/Biopsy Results | | Director: Dr. Carole Bolin<br>4125 Beaumont Road<br>Lansing, MI 48910-8104<br>Phone: 517-353-1683<br>Fax: 517-353-5096<br>www.animalhealth.msu.edu<br>REPORT OF LABORATORY EXAMINATION<br>Client:<br>Virginia Zoological Park (81446)<br>3500 Granby<br>Norfolk, VA 23504<br>Owner:<br>Virginia Zoological/City of Norfolk, -<br>Rcvd Date: 4/28/2011 11:41:00 AM<br>Admitted By: Not, Provided<br>Ordered By: N/A<br>Encounter: 01169614<br>CR#: AP<br>Animal: ELEPHANT MRN:<br>Species: Other Mammal Breed: Elephant<br>Age: 38 years Gender: Female<br>Tag/Reg ID:<br>Other ID:<br>L = Low Result; H = High Result; @ = Critical Result; ^ = Corrected Result; * = Interpretive Data; # = Result Footnote<br>Print Date/Time: 8/4/2011 12:19 PM Page 1 of 7<br>N e c r o p s y P r e l i m i n a r y R e p o r t<br>Accession Number: Received Date/Time: Verified Date/Time: Pathologist:<br>NO-11-0000504 4/28/2011 11:43:00 AM 5/3/2011 8:47:14 AM Agnew, Dalen |

Printed: 6/5/2013 1:18:42 PM Virginia Zoological Park Page 3 Of 24

**Figure S1:** First page of pathology report of the elephant we examined for the anatomical measurements of the elephant.

#### 2 Supplemental Videos

**Supplementary video 1**. Video of elephant lifting 20 kg barbell.

**Supplementary video 2**. Video of elephant lifting 43 kg barbell.

**Supplementary video 3**. Video of elephant lifting 60 kg barbell.

**Supplementary video 4**. Rotating video of CT scan of the Smithsonian elephant trunk, which includes the distal 60 cm.

**Supplementary video 5**. Fly-through video of the CT scan of the Smithsonian elephant trunk. Video starts at a distance 60 from the tip, and each consists of slices that are 0.5mm apart.
